## Supplementary material for "Valuing carbon sequestration by Antarctic krill faecal pellets": All Supplementary Info

Supplementary Information.

Cavan et al.

**Abundance**

KRILLBASE consists of krill density data from net hauls taken throughout the year (Atkinson *et al.*, 2017), such that the time of year sampling occurs could lead to misinterpretation/bias when looking at krill density distribution at a circumpolar scale. For instance, if a grid cell only has net haul data from April, but the adjacent grid cell has data only from January, the first April grid cell would appear to have low krill densities compared to the adjacent January one, when actually this is due to the month the water was sampled for krill. To avoid over- or under-estimating krill density in a particular area, Atkinson et al. (2008) standardised the density data and present circumpolar maps based on when krill density is thought to be highest, December and January. They convert net haul density data to what the density would be on the 1st January after fitting the regressions below with the respective number of days post October 1st, the net mouth area and regression coefficients. See also Table 4 in Atkinson et al. (2017).

*D* = days from October = 1, 32, 62, 93, 124, 152, 183 days
*M* = net mouth area ($m^2$) = 8 $m^2$
*a1* = -0.6478
*b1* = 2.335
*c1* = 0.0204
*d1* = -0.0001086
*a2* = 0.474
*b2* = -0.1912
*c2* = 0.00416
*d2* = -0.00002898

$$G1 = a1 + b1 * log10(M) + c1 * D + d1 * D^2 \quad \text{(Equation S1)}$$

$$G2 = 1 + exp^{\left(a1+b1*log10(M)+c1*D+d1*D^2\right)*10^{\left(1+b2*log10(M)+c2*D+d2*D^2\right)}} \quad \text{(Equation S2)}$$

$$Conversion\ factor = \frac{exp^{\frac{G1}{G2}}}{2.51491418861797} \quad \text{(Equation S3)}$$

This scales up density data to the maximum abundance likely for a particular area or cell in December/January and results in the column in KRILLBASE entitled 'STANDARDISED_ KRILL_UNDER_1M2'. Here in this study, we used this standardised density column and the conversions above to model the abundance in each cell *back* to the 1st day of each month to give estimates of krill density on a circumpolar scale and with time (Fig. S1). Our final estimates of carbon sequestration would be over-estimated if we assumed December/January abundances were relevant throughout the whole Austral spring/summer season. See also 'Abundance continued' section at end of document.

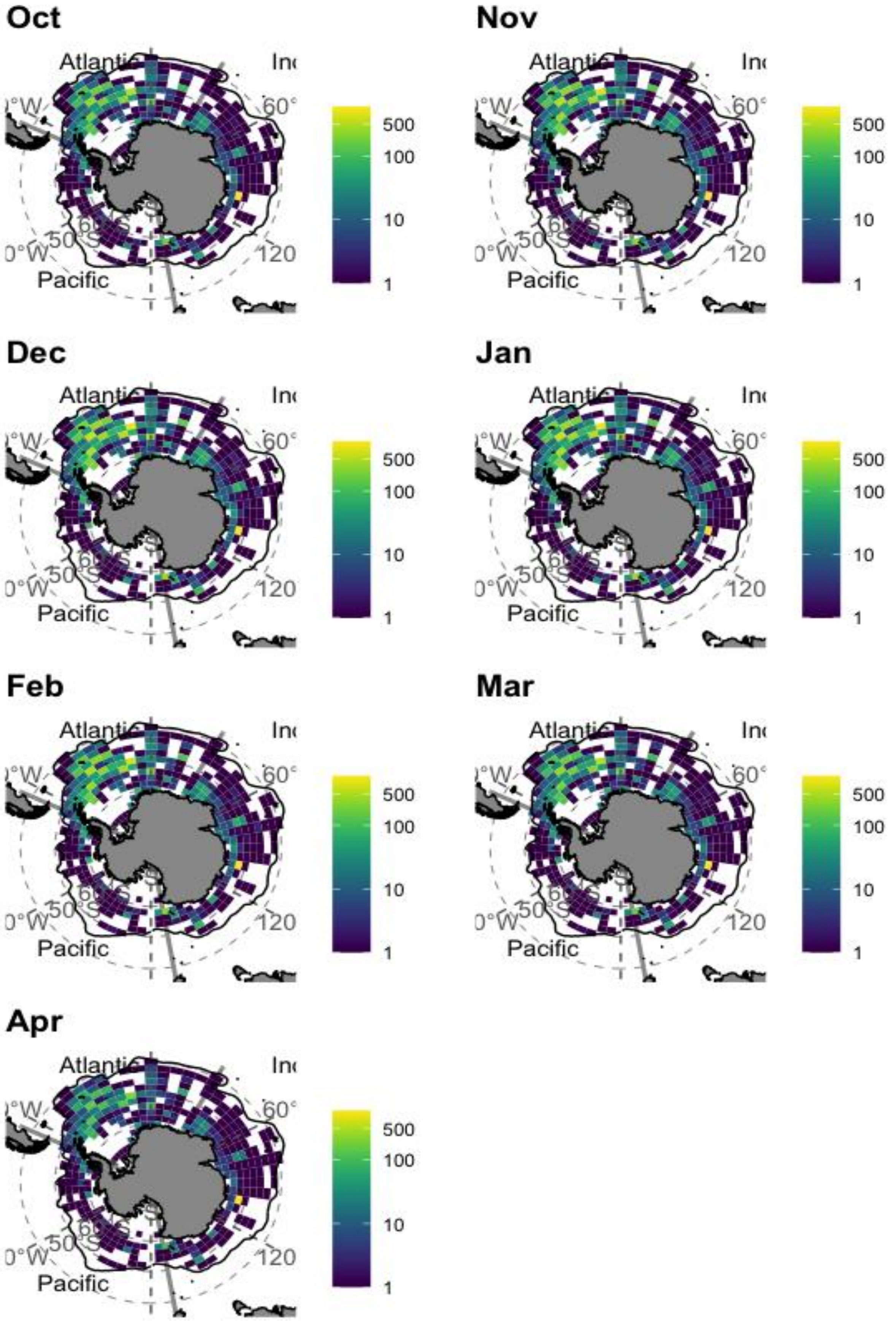

**Supplementary Fig. S1** Krill density (# individuals m$^{-2}$) for each month mapped at 2° x 6° resolution applying the regression model above (Equations S1-3).

**Egestion rate**

Our calculations require a faecal pellet egestion rate (*E*), expressed in mg C per individual per day. Published estimates such as 4.03 mg C $ind^{-1}$ $d^{-1}$ (Clarke *et al.*, 1988) rely on laboratory experiments where food is not limiting to the krill and represent highly productive Spring conditions only. At-sea egestion experiments over Autumn and Spring produce lower egestion rates, from 0.01-0.7 mg C $ind^{-1}$ $d^{-1}$ (Atkinson *et al.*, 2012). Here we explore an appropriate average egestion rate that could be applied across our sampling time series, in the absence of data on how egestion rates vary over time. We use three different approaches to estimate *E* based on 1) circumpolar krill production estimates, 2) daily mass krill both and 3) food-web models. We provide validation for our *E* estimates by converting them to estimates of annual food consumption (in carbon units) by the circumpolar krill stock and comparing this to an indicative estimate of circumpolar primary production (1949 Mt C $y^{-1}$) (Arrigo *et al.*, 2008). Our final median egestion rate of 0.46 mg C $ind^{-1}$ $d^{-1}$, which we use in this study, lies within the range reported by Atkinson et al (2012). See next page.

**Approach 1:** estimates $E$ from published estimates of circumpolar production using the equation:

$$E = \frac{P(1-AE)}{N.GGE.d} \quad \text{(Equation S4)}$$

where $P$ and $N$ are estimates of a circumpolar krill production and abundance respectively, $AE$ is assimilation efficiency (as a fraction), $GGE$ is gross growth efficiency (as a fraction) and $d$ is the number of days in the productive season. We took estimates of $P$ and $N$ from Atkinson *et al.*, (2009). Each of these estimates is specific to an assumed individual mass of krill. We considered small (20mm = 48.4 mg wet mass), typical (40mm = 483 mg wet mass) and large (50 mm = 1127 mg wet mass) krill and converted wet mass to carbon using the conversion factor (0.1075) in Belcher *et al.*, (2019). We also used the $GGE$ and $AE$ values in Belcher *et al.* (2019) and 181 days (representing the 6 months November to April). Results for each set of input values are given in Table S1.

**Table S1:** Assumptions and results of egestion rate calculations using Approach 1. *Q/PP* is circumpolar consumption per unit primary production where consumption ($Q$) is calculated as *P*/*GGE.*

| *P* (Mt C) | *N* (x10$^{14}$) | *GGE* | *AE* | *E* (mg C d$^{-1}$) | *Q/PP* |
|---|---|---|---|---|---|
| **Krill starting size = 20mm (5.2 mg C)** | | | | | |
| 36.77 | 0.79 | 0.20 | 0.42 | 0.75 | 9% |
| 36.77 | 0.79 | 0.20 | 0.75 | 0.32 | 9% |
| 36.77 | 0.79 | 0.20 | 0.85 | 0.19 | 9% |
| 36.77 | 0.79 | 0.20 | 0.94 | 0.08 | 9% |
| 36.77 | 0.79 | 0.30 | 0.42 | 0.50 | 6% |
| 36.77 | 0.79 | 0.30 | 0.75 | 0.22 | 6% |
| 36.77 | 0.79 | 0.30 | 0.85 | 0.13 | 6% |
| 36.77 | 0.79 | 0.30 | 0.94 | 0.05 | 6% |
| **mean** | | | | 0.28 | 8% |
| **Krill starting size = 40mm (40.53 mg C)** | | | | | |
| 52.68 | 0.78 | 0.20 | 0.42 | 1.08 | 14% |
| 52.68 | 0.78 | 0.20 | 0.75 | 0.47 | 14% |
| 52.68 | 0.78 | 0.20 | 0.85 | 0.28 | 14% |
| 52.68 | 0.78 | 0.20 | 0.94 | 0.11 | 14% |
| 52.68 | 0.78 | 0.30 | 0.42 | 0.72 | 9% |
| 52.68 | 0.78 | 0.30 | 0.75 | 0.31 | 9% |
| 52.68 | 0.78 | 0.30 | 0.85 | 0.19 | 9% |
| 52.68 | 0.78 | 0.30 | 0.94 | 0.07 | 9% |
| **mean** | | | | 0.40 | 11% |
| **Krill starting size = 50mm (94.49 mg C)** | | | | | |
| 57.62 | 0.78 | 0.20 | 0.42 | 1.18 | 15% |
| 57.62 | 0.78 | 0.20 | 0.75 | 0.51 | 15% |
| 57.62 | 0.78 | 0.20 | 0.85 | 0.31 | 15% |
| 57.62 | 0.78 | 0.20 | 0.94 | 0.12 | 15% |

| | | | | | |
|---|---|---|---|---|---|
| 57.62 | 0.78 | 0.30 | 0.42 | 0.79 | 10% |
| 57.62 | 0.78 | 0.30 | 0.75 | 0.34 | 10% |
| 57.62 | 0.78 | 0.30 | 0.85 | 0.20 | 10% |
| **mean** | | | | 0.48 | 10% |

**Approach 2**: calculates the daily egestion rate of individual krill based on observed daily mass growth:

$$E = \frac{(1-AE).m.GR}{GGE} \qquad \text{(Equation S5)}$$

where *m* is individual mean mass at the start of the growth season in mg C and *GR* is daily growth rate as a proportion of *m*. We used a *GR* estimate of 1.17% (Atkinson *et al.*, 2006) alongside the values for other parameters assumed in Approach 1. Results for each set of input values are given in Table S2.

**Table S2:** Assumptions and results of egestion rate calculations using Approach 2. *Q/PP* is circumpolar consumption per unit primary production where consumption (*Q*) is calculated as *m.GR.d* /(*GGE.NI*) where *N* is taken from Table S1 for the relevant krill size and *d* is 181.

| *GR* (%) | *m* (mg C) | *GGE* | *AE* | *E* (mg C $d^{-1}$ | *Q/PP* |
|---|---|---|---|---|---|
| **Krill starting size = 20mm (5.2 mg C)** | | | | | |
| 1.17 | 5.20 | 0.20 | 0.42 | 0.18 | 2% |
| 1.17 | 5.20 | 0.20 | 0.75 | 0.08 | 2% |
| 1.17 | 5.20 | 0.20 | 0.85 | 0.05 | 2% |
| 1.17 | 5.20 | 0.20 | 0.94 | 0.02 | 2% |
| 1.17 | 5.20 | 0.30 | 0.42 | 0.12 | 1% |
| 1.17 | 5.20 | 0.30 | 0.75 | 0.05 | 1% |
| 1.17 | 5.20 | 0.30 | 0.85 | 0.03 | 1% |
| 1.17 | 5.20 | 0.30 | 0.94 | 0.01 | 1% |
| **mean** | | | | 0.07 | 2% |
| **Krill starting size = 40mm (40.53 mg C)** | | | | | |
| 0.01 | 51.92 | 0.20 | 0.42 | 1.76 | 22% |
| 0.01 | 51.92 | 0.20 | 0.75 | 0.76 | 22% |
| 0.01 | 51.92 | 0.20 | 0.85 | 0.46 | 22% |
| 0.01 | 51.92 | 0.20 | 0.94 | 0.18 | 22% |
| 0.01 | 51.92 | 0.30 | 0.42 | 1.17 | 15% |
| 0.01 | 51.92 | 0.30 | 0.75 | 0.51 | 15% |
| 0.01 | 51.92 | 0.30 | 0.85 | 0.30 | 15% |
| 0.01 | 51.92 | 0.30 | 0.94 | 0.12 | 15% |
| **mean** | | | | 0.66 | 18% |
| **Krill starting size = 50mm (94.49 mg C)** | | | | | |
| 0.01 | 121.15 | 0.20 | 0.42 | 4.11 | 51% |
| 0.01 | 121.15 | 0.20 | 0.75 | 1.77 | 51% |
| 0.01 | 121.15 | 0.20 | 0.85 | 1.06 | 51% |
| 0.01 | 121.15 | 0.20 | 0.94 | 0.43 | 51% |
| 0.01 | 121.15 | 0.30 | 0.42 | 2.74 | 34% |
| 0.01 | 121.15 | 0.30 | 0.75 | 1.18 | 34% |
| 0.01 | 121.15 | 0.30 | 0.85 | 0.71 | 34% |
| 0.01 | 121.15 | 0.30 | 0.94 | 0.28 | 34% |
| **mean** | | | | 1.54 | 43% |

**Approach 3:** uses parameters from three regional foodweb models for habitats south of the Antarctic Polar Front compiled by Hill *et al.* (2021).

$$E = \frac{(1-AE).\frac{Q}{B}.B}{\sigma.d} \qquad \text{(Equation S6)}$$

Where $\frac{Q}{B}$ is the annual carbon consumption per unit krill carbon biomass, *B*, per unit model area; σ is the number of individuals per unit model area, calculated from *B* and an assumed individual mean mass, *m*, and *d* is 181. Results for each set of input values are given in Table S3.

**Table S3:** Assumptions and results of egestion rate calculations using Approach 3. Model is the specific ecosystem model reported in Hill *et al.*, (2021) (see their Appendix A). *Q/PP* is circumpolar consumption per unit primary production where consumption (*Q*) is calculated as *N.Q/B.m* where *N* is taken from Table S1 for the relevant krill size and *m* is taken from Table S2 for the relevant krill size.

| *Model* | *Q/B* | *B (kg m$^{-2}$)* | *AE* | *E* (mg C d$^{-1}$) | *Q/PP* |
|---|---|---|---|---|---|
| **Krill size = 20mm (5.2 mg C)** | | | | | |
| SG-Aggr | 16.00 | 2.55 | 0.80 | 0.09 | 3% |
| AP-Aggr | 4.70 | 2.34 | 0.73 | 0.04 | 1% |
| RS-Aggr | 10.10 | 0.14 | 0.80 | 0.06 | 2% |
| **mean** | | | | 0.06 | 2% |
| **Krill starting size = 40mm (40.53 mg C)** | | | | | |
| SG-Aggr | 16.00 | 2.55 | 0.80 | 0.92 | 33% |
| AP-Aggr | 4.70 | 2.34 | 0.73 | 0.37 | 10% |
| RS-Aggr | 10.10 | 0.14 | 0.80 | 0.58 | 21% |
| **mean** | | | | 0.62 | 21% |
| **Krill starting size = 50mm (94.49 mg C** | | | | | |
| SG-Aggr | 16.00 | 2.55 | 0.80 | 2.14 | 78% |
| AP-Aggr | 4.70 | 2.34 | 0.73 | 0.86 | 23% |
| RS-Aggr | 10.10 | 0.14 | 0.80 | 1.35 | 49% |
| **mean** | | | | 1.45 | 50% |

These three approaches, used with various parameter combinations, give a range of individual egestion rates spanning three orders of magnitude from 0.01 to 4.11 mg C $ind^{-1}$ $d^{-1}$. Values greater than 1 mg C $ind^{-1}$ $d^{-1}$ occur only when either the extreme low value (0.42) is used for *AE* or krill size is assumed to be 50 mm. Values less than 0.1 mg C $ind^{-1}$ $d^{-1}$ occur only when either the extreme high value (0.94) is used for *AE* or krill size is assumed to be 20 mm.

There is incomplete overlap between the habitat of Antarctic krill (waters south of the Antarctic Polar Front; Atkinson et al (2008)) and the area that the circumpolar primary production estimate applies to (waters south of 50°S; (Arrigo *et al.*, 2008)). The comparison is therefore indicative only. Consumption per unit primary production (*Q/PP*) values >30% are unlikely given that krill constitutes approximately 30% of metazoan grazer biomass in the Southern Ocean (Yang *et al.*, 2022) and consumption by metazoans is only one of several possible fates of primary production.  This comparison demonstrates that the assumption of low *AE* and/or large average krill size can lead to implausible estimates of circumpolar consumption and therefore egestion rate.

A large circumpolar database of postlarval krill lengths suggests that an appropriate average length is ~ 40mm (Atkinson *et al.*, 2009). Thus we use the median of egestion rate estimates for 40 mm krill (0.46 mg C $d^{-1}$) in our main analysis, and conduct a sensitivity analysis in the main text using the 5$^{th}$ and 95$^{th}$ percentiles of estimates for 40 mm krill (0.11 and 1.23 mg C $d^{-1}$ respectively). Our egestion estimates are considerably lower than some values used in previous studies (e.g. Clarke et al (1988) used a rate of 4.03 mg C $d^{-1}$ and Belcher *et al.*, (2019) used a rate of 3.2 mg C $d^{-1}$). These values were based on egestion rates observed over a period of one hour which were then multiplied by 24 to give daily rates (Clarke *et al.*, 1988). Our calculations suggest that these rates are not likely to be sustained throughout the summer season or at the circumpolar scale.

### Martins b analysis

**Table S4: Martin's b for krill faecal pellet POC flux adapted from Belcher et al. 2017.** The median *b* is -0.30, a slight change from Belcher et al. of -0.32 due to the addition of Pauli *et al.*, (2021) data. We use *b* = -0.3 in this study.

| Source | Region | Depth (m) | Season | Krill FP flux (mg C $m^{-2}$ $d^{-1}$) | Attenuation (*b*) |
|---|---|---|---|---|---|
| Belcher et al. (2017)(Belcher *et al.*, 2017) | South Orkneys | 64 | December | 66.7 | 0.13 |
| | | 165 | | 75.5 | |
| | | 76 | December | 33.0 | 1.8 |
| | | 178 | | 154.1 | |
| | | 61 | November | 68.0 | 0.13 |
| | | 163 | | 77.3 | |
| | | 150 | | *205* | |
| Wefer at al. (1988)(Wefer *et al.*, 1988) [b] | Bransfield Strait | 494 | January | 281.2 | -0.6 |
| | | 1588 | | 139.9 | |
| Accornero et al. (2003)(Accornero *et al.*, 2003) | Ross sea | 180 | Annual mean | 0.05 | -0.32 |
| | | 868 | | 0.03 | |
| Cavan et al. (2015)(Cavan *et al.*, 2015) [c] | Scotia Sea | 70 | January | 58.6 | 0.32 |
| | | 170 | | 77.9 | |
| González (1992)(González, 1992) [d] | Scotia-Weddell seas | 50 | December-January | 10 | -0.63 |
| | | 150 | | 5 | |
| | | 50 | December-January | 5.5 | -2.2 |
| | | 150 | | 0.5 | |
| | | 50 | December-January | 3 | 0.66 |
| | | 150 | | 10.5 | |
| | | 300 | | 9 | |
| | | 50 | December-January | 22.5 | -2.5 |
| | | 150 | | 1.5 | |
| Pauli et al. (2021) | Elephant Island | 100 | April | 35.05 | -0.61 |
| | | 200 | | 8.47 | |
| | | 300 | | 21.69 | |
| | | 100 | April | 16.19 | 0.98 |
| | | 200 | | 17.7 | |
| | | 300 | | 53.24 | |
| | | 100 | April | 25.26 | -0.28 |
| | | 200 | | 11.68 | |
| | | 300 | | 20.62 | |
| | | 100 | April | 6.76 | 0.44 |
| | | 200 | | 50.23 | |
| | | 300 | | 13.80 | |
| | | 100 | April | 28.85 | -0.23 |
| | | 200 | | 42.36 | |
| | | 300 | | 20.16 | |

[b] Fluxes are for total particulate organic carbon
[c] Fluxes are for all FP, but krill FP were dominant
[d] Fluxes are FP in terms of FP dry weight, and have been estimated from Fig. 3, Fig. 5 of González (1992)(González, 1992)

**Table S5 Sensitivity Analysis, see Fig. 3 in main text**. Using original model parameters the total carbon sequestered from krill faeces is 19.5 MtC. Increasing krill density (abundance), egestion rate and attenuation rate of sinking pellet POC (more positive) increases MtC sequestered, whilst increasing sequestration depth decreases total MtC sequestered. Where means are given these refer to mean across whole time series, October to April and are presented in the Table for comparison across analyses, but individual grid cell values are used to calculate the new MtC values.

| | | | **Sensitivity Analysis** | | | **Parameter uncertainty** | | |
|---|---|---|---|---|---|---|---|---|
| **Parameter** | **Symbol** | **Original value** | **New value (+/- 10 %)** | **New MtC** | **Percentage change** | **New value** | **New MtC** | **Percentage change** |
| **Increase in parameter value** | | | | | | | | |
| Density | *N* | Mean = 20 ind. $m^{-2}$ | Mean = 22 ind. $m^{-2}$ | 21.6 | 110 % | Mean = 36 ind $m^{-2}$ | 35.5 | 180 % |
| Egestion | *E* | 0.46 mg C $d^{-1}$ | 0.51 mg C $d^{-1}$ | 21.6 | 110 % | 1.23 mg C $d^{-1}$ | 52.5 | 267 % |
| Sequestration depth | $FPT_{100}$ | Mean = 381 m | Mean = 419 m | 19.1 | 97 % | Mean = 619 m | 17.0 | 87 % |
| Attenuation rate | *b* | -0.3 | -0.27 | 21.1 | 108 % | +0.13 | 62.8 | 320 % |
| **Decrease in parameter value** | | | | | | | | |
| Density | *N* | Mean = 20 ind. $m^{-2}$ | Mean =18 ind $m^{-2}$ | 17.7 | 90 % | Mean = 4 ind. $m^{-2}$ | 3.9 | 20 % |
| Egestion | *E* | 0.46 mg C $d^{-1}$ | 0.41 mg C $d^{-1}$ | 17.7 | 90 % | 0.11 mg C $d^{-1}$ | 4.7 | 24 % |
| Sequestration depth | $FPT_{100}$ | Mean = 381 m | Mean = 343 m | 20.0 | 103 % | Mean = 187 m | 24.2 | 123 % |
| Attenuation rate | *b* | -0.3 | -0.33 | 18.1 | 92 % | -0.61 | 8.5 | 44 % |

### Validating our model

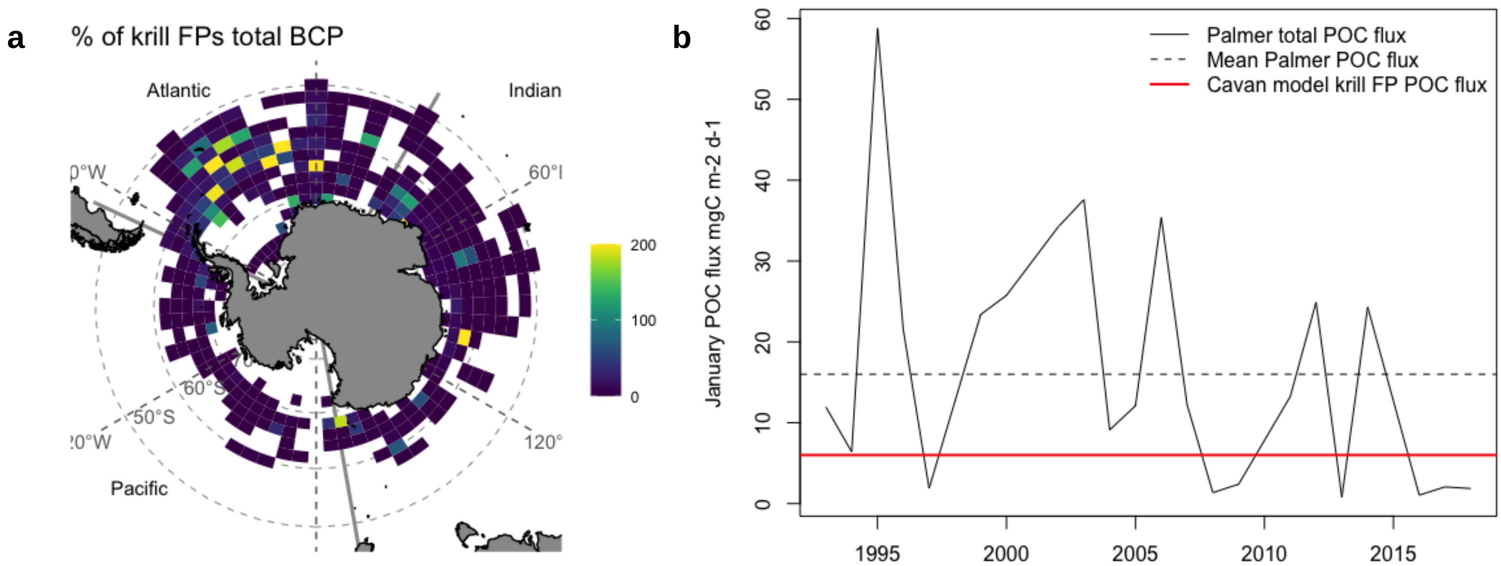

**Supplementary Fig S2. Krill FP sense check.** a) shows the percentage contribution of krill faecal pellets to total POC flux at the sequestration depth, with the total POC flux from copepods and plankton predicted by the DeVries and Weber (2017) model. In some locations when krill are abundant high krill POC flux (yellow pixels) suggest the DeVries and Weber model underestimates fluxes in some regions, as like most biogeochemical models it does not include micronekton or krill. b) shows the January sediment trap time-series data at Palmer station (Trinh *et al.*, 2023) and the average total POC flux of their January time series (dashed black line) at 170 m depth. The red line shows the average from our model of adult krill FP POC flux only, which here near the Antarctic continent represents ~ 38 % of the total flux.

### Transport matrix

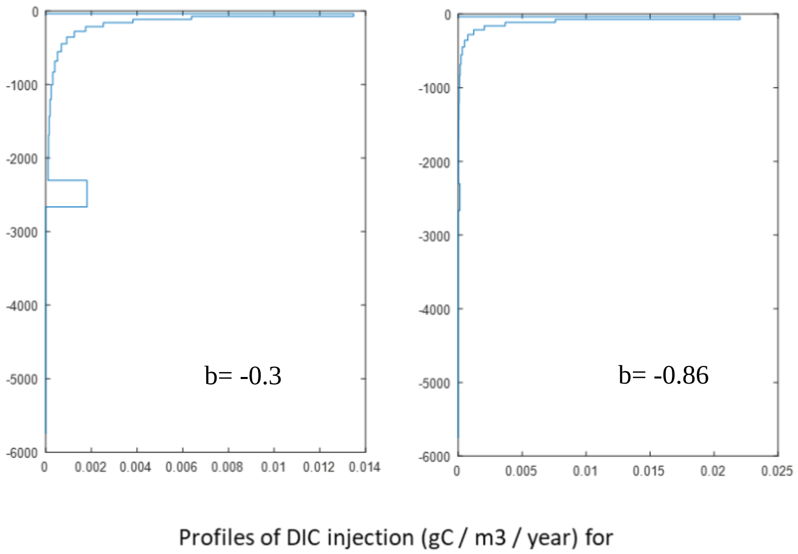

**Supplementary FigS3**. Fraction of pellet carbon attenuating with depth, for when Martin's b is set to -0.3 to represent observations of krill faecal pellets, or the Martin *et al.*, (1987) value from the equatorial Pacific, of -0.86. These data feed into the OCIM transport matrix to determine the fate of pellet-originating carbon. See Fig. 4 in the main text for when b = -0.30, and Fig. S4 below for when b = -0.86.

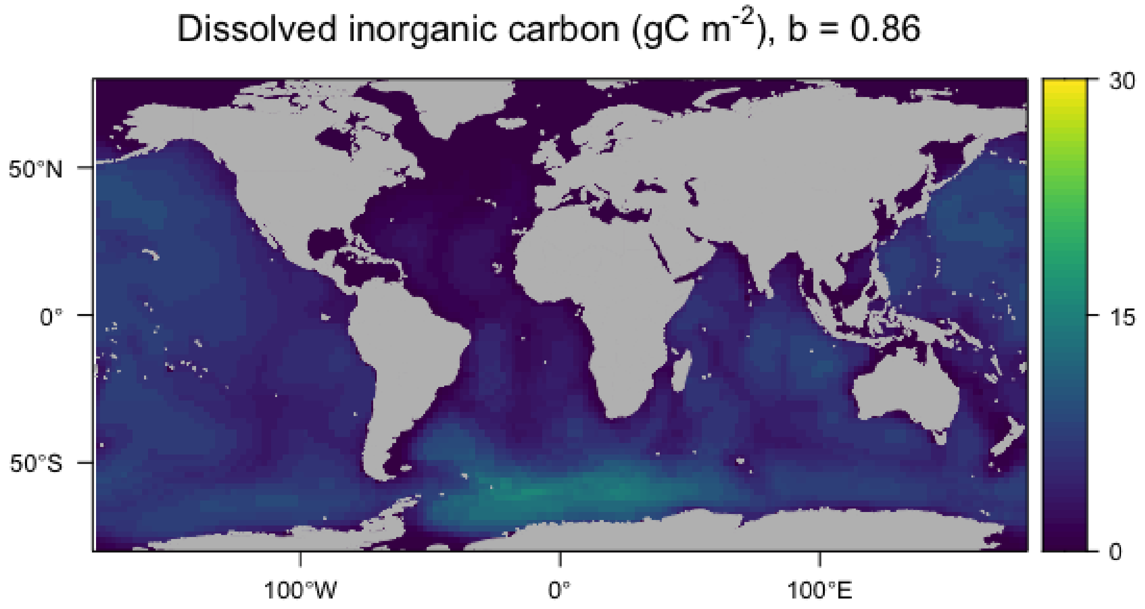

**Supplementary Fig S4**. Concentration of DIC (gC m$^{-2}$) through the water column when attenuation is set to -0.86. The resulting carbon stored equates to 1.7 PgC for an average of 58 years.

**Moult, carcasses and migrations**

To gauge the total carbon sequestration krill may have in addition to their faecal pellet, we estimate the magnitude of other contributions from krill moults, carcasses, and vertical migrations. Moults can sink at similar rates to pellets, with sinking rates ranging from 50-1000 m $d^{-1}$ (Nicol and Stolp, 1989) compared to pellets which range from 27–1218 m $d^{-1}$(Atkinson *et al.*, 2012). The organic carbon content of moults is also high, ~ 73 % of dry mass (Nicol and Stolp, 1989). A sediment trap near South Georgia provides monthly krill moult estimates, which throughout krill productive months (the temporal scale used in this study) equals the carbon fluxes from krill faecal pellets (Manno *et al.*, 2020). Therefore in Fig. 5 in the main text we suggest the magnitude of carbon fluxes from krill moults would be equal to krill pellets. For carcasses, Manno et al. 2020 show the contribution is more variable and highest in winter when krill mortality peaks. As the winter months are not included in our analysis, and due to the more limited data and knowledge on krill carcass contributions to sinking flux, we chose not to report circumpolar estimates in flux for carcasses.

Daily and seasonal migrations of krill can actively transfer $CO_2$ into the mesopelagic zone and act as efficient vectors of carbon. Total krill biomass is 380 Mt wet mass (Atkinson 2009), of which about 10 % is carbon (38 MtC), with 87 % of this carbon mass in the open ocean and the remaining 13 % living on the shelf (Atkinson *et al.*, 2008). Estimates from Schmidt *et al.*, (2011) suggest ~19 % of krill in the open ocean and 2 % of krill living on the shelf reside at 400 m depth at any given time, with these krill undergoing seasonal and diel vertical migrations. Applying these numbers to the krill population would mean ~6 Mt and 0.1 Mt of krill biomass are living deeper than 400 m in the open ocean and on the coast, respectively. We assume krill respire 2 % of their biomass in terms of carbon each day, based on estimates of 1 % for egestion, 1 % for growth and if 4 % is available then 2 % remains for respiration to yield a gross growth efficiency of 25 % (Atkinson *et al.*, 2006). This yields a total carbon release from respiration at 400 m of 47 Mt C. Both the fraction of krill living below 400 m and the growth gross efficiencies are based on summer values, and so we recalculate the total respiration leaving it constant from December to March, but reduce it to a third of the value for the remainder of the year. This gives a more conservative and more likely realistic value of 26 MtC released by krill. A recent modelling study quantified the contributions of deep metazoan respiration to carbon sequestration on a large spatial scale across the sub-polar to tropical global oceans. They estimated non-polar biomass of macrozooplankton to be 80 Mt C, and the respired carbon released to be 50 Mt C, suggesting macrozooplankton respire 63 % of their total biomass (Pinti *et al.*, 2023). Our estimates are similar, and if krill biomass is 38 MtC and they respire 26 Mt C, this is 68 % of the total biomass. Note krill biomasses are total mass and not just carbon mass.

We repeat the analysis based on the circumpolar abundances calculated in this study, which peak at 5.7 x $10^{14}$ individual krill. Using the abundances in this study we can convert the monthly krill density data using the conversion factors in Atkinson *et al.*, (2009) (see Table 4) to monthly biomass. With monthly biomass we use the same fractions of open ocean krill and those living below 400 m, and a 2 % respiration rate and find that for our 7-month time series the carbon respired is 14 Mt C, and 70 % of the total average biomass (22 Mt). Using the lowest monthly value for carbon respiration (due to lower biomass) of April (1.4 MtC), we assume the remaining months of the year, May through September, have the same respiration levels. Including these brings the total annual carbon sequestered through deep respiration to 21 MtC, similar to the other value we calculated of 26 MtC with the difference due to densities and how the circumpolar grid has been sampled spatially.

**Abundance continued**

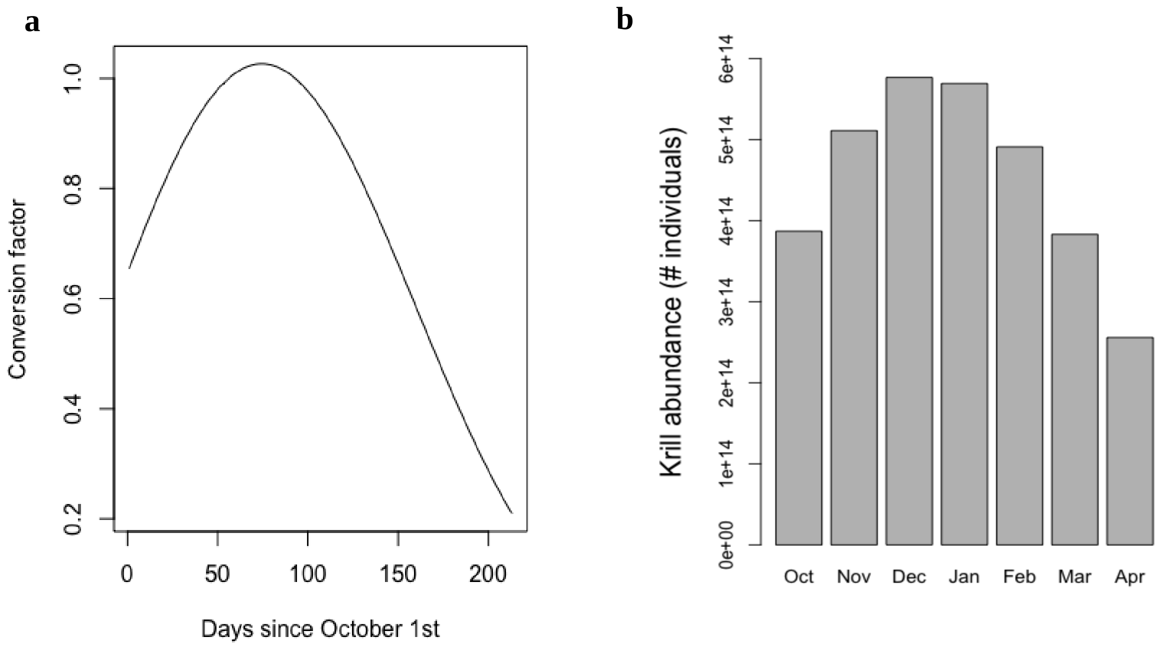

**Supplementary Fig. S5** Conversion factors applied to krill density (a), with a conversion of 1 in January, and slightly >1 in December. This results in circumpolar abundance estimates of krill each month (b) which are highest in December and January, and lowest in April. Note here the abundance is presented after the extreme values of > 600 ind $m^{-2}$ are capped at 600 ind $m^{-2}$.

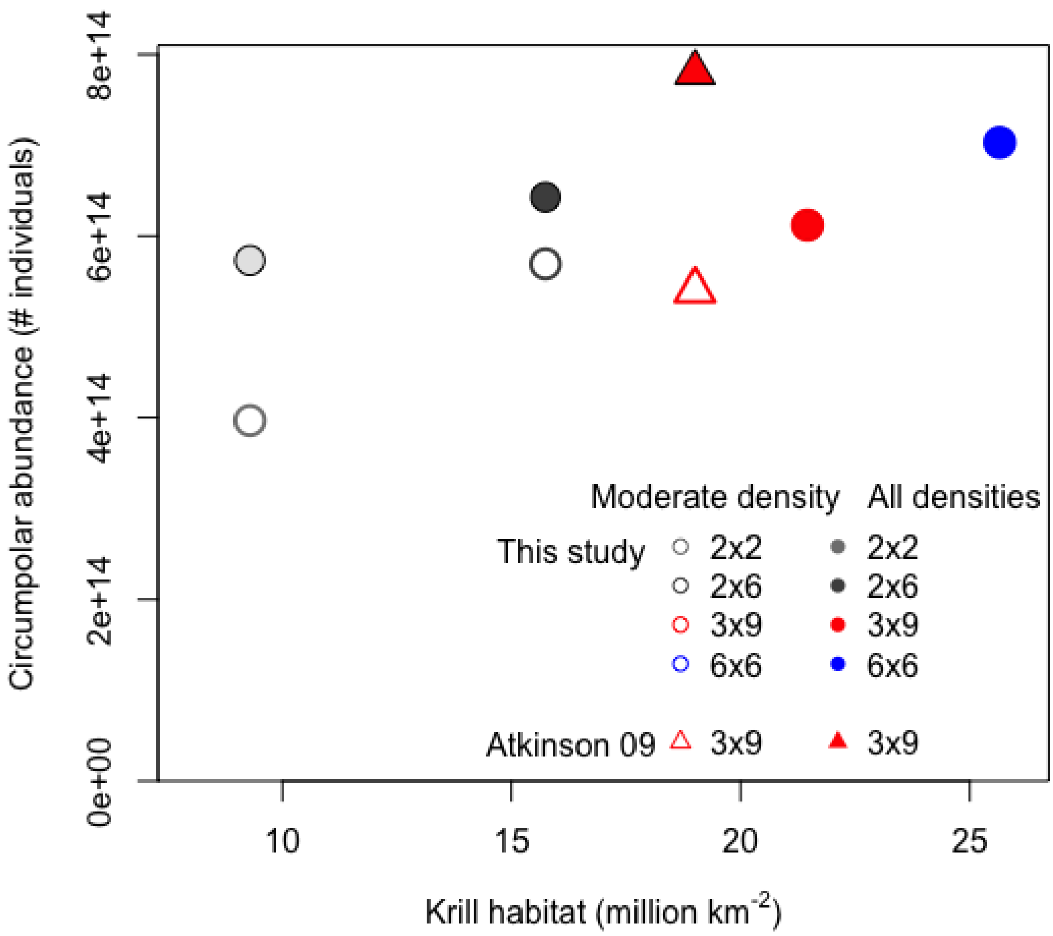

**Supplementary Fig. S6.** Change in krill density with resolution and size of area sampled. Light grey point is at a 2°x2° resolution, dark grey is 2°x6° resolution used in this study, red point is 3°x9° resolution and blue is 6°x 6° resolution. The closed points refer to krill abundance as presented in KRILLBASE, and the open points refer to krill abundance data as in KRILLBASE however with densities >600 ind $m^{-2}$ capped at 600 ind $m^{-2}$. This excludes bias from a few extremely high net catches sampling in a swarm, but still allows for high krill densities to occur (mean = 28 ind $m^{-2}$ within the defined krill habitat, i.e. krill density > 0 ind $m^{-2}$, and 20 ind $m^{-2}$ when including 0 ind $m^{-2}$ grid cells). Open points for the 3°x9° resolution (red) and 6°x 6° resolution (blue) are the same as the closed points, as averaging over wider areas of ocean lowered the mean individual krill per $m^{-2}$, so that none were > 600 $m^{-2}$. These resolutions do increase the sampling area (Fig. S7) which directly impacts total carbon sequestered and hence we use a 2°x6° resolution. The closed red triangle is the abundance measured at 3°x9° resolution by Atkinson *et al.*, (2009) using all KRILLBASE data, and the open triangle using their moderate values, where they reduced every sample >300 ind $m^{-2}$ to their mean of 36 ind $m^{-2}$. The discrepancies between the closed red point and the closed red triangle in terms of krill habitat (x-axis) are likely due to different gridding approaches between the Atkinson 2009 paper and our study. The final circumpolar abundance we use in this study is $5.7e^{14}$ at a resolution of 2°x6° (dark grey open point), similar to the final abundance of $5.4e^{14}$ from Atkinson 2009 (open red triangle).

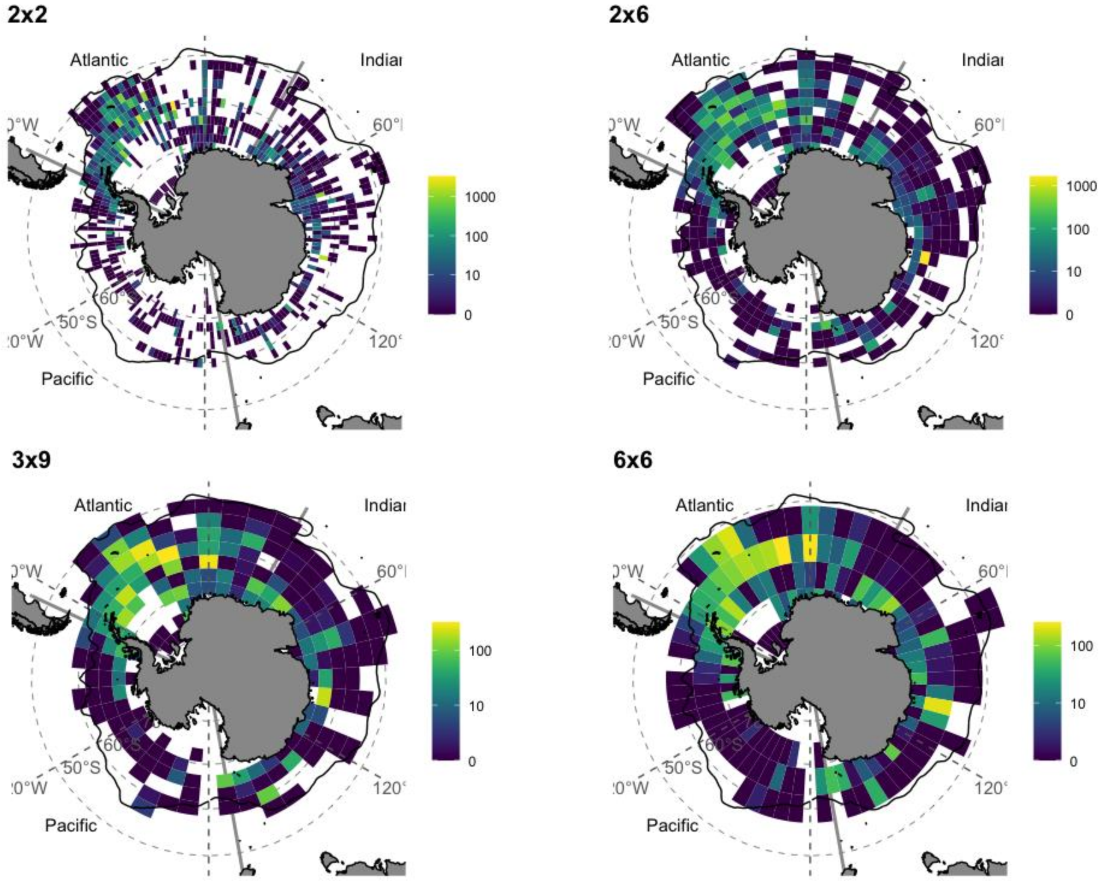

**Fig. S7**. Circumpolar densities of krill at different resolutions. Here the high densities have been left (i.e. not capped at 600 ind m$^{-2}$), to show that reducing the resolution reduces the high mean abundance values in each grid cell, whilst increases the habitat area. This is why you do not see a linear trend in krill habitat with abundance in Fig. S6, because although a large area is sampled, the mean densities in each grid are lower.
